# Ketone Bodies in Right Ventricular Failure: A Unique Therapeutic Opportunity

**DOI:** 10.1101/2023.04.26.538410

**Authors:** Madelyn Blake, Patrycja Puchalska, Felipe Kazmirczak, Thenappan Thenappan, Peter A. Crawford, Kurt W. Prins

## Abstract

Ketone bodies are pleotropic metabolites that play important roles in multiple biological processes ranging from bioenergetics to inflammation regulation via suppression of the NLRP3 inflammasome, and epigenetic modifications. Ketone bodies are elevated in left ventricular failure (LVF) and multiple approaches that increase ketone concentrations exert advantageous cardiac effects in rodents and humans. However, the relationships between ketone bodies and right ventricular failure (RVF) are relatively unexplored. Moreover, the cardioprotective properties of ketones in preclinical RVF are unknown. Here, we show a compensatory ketosis is absent in pulmonary arterial hypertension (PAH) patients with RVF. In the monocrotaline (MCT) rat model of PAH-mediated RVF, a dietary-induced ketosis improves RV function, suppresses NLRP3 inflammasome activation, and combats RV fibrosis. The summation of these data suggest ketogenic therapies may be particularly efficacious in RVF, and therefore future studies evaluating ketogenic interventions in human RVF are warranted.

## Main

RVF is the leading cause of death in pulmonary arterial hypertension (PAH), but at present there are no therapies that effectively combat RVF (1). This is in juxtaposition to LVF treatment as several medications can both improve LV function and long-term survival (2). Additionally, there are differing degrees of end-organ dysfunction when comparing RVF and LVF. In particular, RVF is associated with more severe liver impairments, which can ultimately result in cirrhosis (3). This raises the possibility that a RV-liver axis exists, and it could serve as a novel therapeutic target for this uniformly lethal consequence of PAH.

Ketone bodies, metabolites that are primarily synthesized by the liver, regulate diverse biological functions ranging from energy homeostasis, inflammation, and epigenetic regulation (4). Both preclinical models and patients with LVF have elevated ketone levels, and ketone concentrations are inversely associated with echocardiographic and biochemical measures of LV function (5). Multiple interventions to induce ketogenic states including intermittent fasting, ketogenic diets, ketone body injections, sodium-glucose co-transporter-2 (SGLT2) inhibitor use, and genetic manipulation of ketone metabolism impart cardio-protective changes in both preclinical and human LVF (6,7). At present, most of the studies evaluating the advantageous effects of ketone bodies in cardiac dysfunction have focused on the favorable metabolic properties that ketone bodies possess (6,7). However, the ketone body β-hydroxybutyrate (βOHB) suppresses nucleotide-binding domain, leucine-rich-containing family, pyrin domain-containing-3 (NLRP3) inflammasome activity (8), which suggests ketogenic interventions may have important anti-inflammatory ramifications that could also combat cardiac dysfunction. We recently showed the NLRP3 inflammasome pathway promotes PAH-mediated RVF (9), but the interplay between RV inflammation/NLRP3 activation, RV function, and circulating ketones has not been studied in-depth.

To address these important knowledge gaps, we performed a translational study that first evaluated relationships between RV function and serum ketone body levels in 51 patients with PAH. Then, we probed the effects of a ketogenic diet on RV function, NLRP3 inflammasome activation, and RV fibrosis in preclinical RVF. We showed human RVF patients lack a compensatory ketosis as serum ketone body concentrations were not associated with hemodynamic, biochemical, or echocardiographic measures of RV function. In rodent studies, a ketogenic diet increased circulating ketones, suppressed NLRP3 inflammasome activation, blunted RV fibrosis, and augmented RV function. In total, our preclinical and clinical data suggest a therapeutic ketosis could combat RVF. Moreover, patients with RVF may be uniquely primed for ketogenic interventions as they exhibit an insufficient ketogenic response.

## Results

### Circulating Ketone Bodies Were Not Strongly Associated with Biochemical, Echocardiographic, or Hemodynamic Measures of RV Function in Human PAH-Mediated RVF

We first validated divergent RV phenotypes when we dichotomized our cohort into two groups: patients with either compensated or decompensated RV function. Hemodynamic, echocardiographic, and biochemical markers of RV function were more severely deranged in the decompensated group (**Table 1 and Supplemental Figure 1**). When comparing clinical characteristics between the two populations, there were no significant differences in cardiovascular comorbidities, age, and sex distributions. However, laboratory assessments revealed higher serum NT pro-BNP and total bilirubin in the decompensated group (**Table 1**), which suggested liver dysfunction was present when RV function was compromised. Finally, the decompensated patients had more advanced PAH with higher mean pulmonary arterial pressure (mPAP) and pulmonary vascular resistance (PVR) (**Table 1**).

**Table 1:** Clinical Characterization of PAH Patients Evaluated in Study.

| **Table 1 Clinical, Echocardiographic, and Hemodynamic Characteristics of Study Cohort** |  |  |  |  |
| --- | --- | --- | --- | --- |
| Characteristics | Total Cohort<br>(n = 51) | Preserved RV Function<br>(n = 26) | Depressed RV Function<br>(n = 25) | p-value |
| Age, years | 58 ± 15 | 59 ± 12 | 57 ± 17 | 0.56 |
| Female, n (%) | 31 (60) | 17 (65) | 14 (56) | 0.62 |
| Body mass index, kg/m<sup>2</sup> | 28 ± 7 | 28 ± 7 | 28 ± 6 | 0.61 |
| WHO Functional Class (n = 48) |  |  |  |  |
| I | 2 (4) | 2 (9) | 0 (0) |  |
| II | 13 (27) | 8 (35) | 5 (20) |  |
| III | 32 (67) | 13 (57) | 19 (76) |  |
| IV | 1 (2) | 0 (0) | 1 (4) |  |
| Comorbidities, n (%) |  |  |  |  |
| Hypertension | 23 (45) | 12 (46) | 11 (44) | 0.76 |
| Diabetes | 12 (24) | 6 (23) | 6 (24) | 0.85 |
| Hyperlipidemia | 16 (31) | 9 (35) | 7 (28) | 0.69 |
| Coronary artery disease | 7 (14) | 3 (12) | 4 (16) | 0.92 |
| Atrial fibrillation | 5 (10) | 1 (4) | 4 (16) | 0.14 |
| COPD | 8 (16) | 5 (19) | 3 (12) | 0.28 |
| Medications, n (%) (n = 49) |  |  |  |  |
| Monotherapy | 7 (14) | 4 (15) | 3 (13) | 0.77 |
| Dual oral therapy | 15 (31) | 12 (46) | 3 (13) | < 0.01 |
| Triple oral therapy | 12 (24) | 5 (19) | 7 (30) | 0.63 |
| Parental Prostacyclin plus oral | 15 (31) | 5 (19) | 10 (43) | 0.09 |
| Lab |  |  |  |  |
| Serum hemoglobin, g/dl (n=51) | 13 ± 2 | 13 ± 2 | 14 ± 2 | 0.34 |
| Serum creatinine, mg/dl (n=51) | 1 ± 0.40 | 1 ± 0.27 | 1 ± 0.43 | 0.36 |
| Serum NT-proBNP (n=50) | 1786 ± 2705 | 508 ± 1067 | 2988 ± 3220 | < 0.001 |
| Serum Bilirubin | 0.67 ± 0.33 | 0.56 ± 0.23 | 0.76 ± 0.39 | < 0.05 |
| Serum ALT | 26 ± 10 | 28 ± 12 | 24 ± 8 | 0.30 |
| Serum AST | 21 ± 9 | 20 ± 8 | 23 ± 9 | 0.21 |
| Echocardiography |  |  |  |  |
| Right ventricular FAC (n=44) | 33 ± 10 | 39 ± 7 | 26 ± 8 | < 0.001 |
| RVEDA | 26 ± 8 | 24 ± 7 | 29 ± 9 | < 0.05 |
| RVESA | 18 ± 8 | 15 ± 6 | 22 ± 9 | < 0.01 |
| TAPSE | 2 ± 0.63 | 2.4 ± 0.64 | 1.6 ± 0.38 | < 0.001 |
| S' | 11 ± 4 | 13 ± 4 | 9 ± 3 | < 0.01 |
| Tricuspid regurgitation severity | 3 ± 1 | 2 ± 1 | 3 ± 1 | < 0.05 |
| Hemodynamics |  |  |  |  |
| Heart rate, beats/min (n=51) | 74 ± 19 | 71 ± 12 | 77 ± 24 | 0.25 |
| Mean right atrial, mm Hg (n =51) | 7 ± 5 | 7 ± 4 | 11 ± 6 | < 0.005 |
| Mean PAP, mm Hg (n=51) | 43 ± 12 | 36 ± 8 | 49 ± 14 | < 0.001 |
| Cardiac output, liters/min (n=51) | 5 ± 2 | 6 ± 2 | 4 ± 1 | < 0.001 |
| Cardiac index, liters/min/m<sup>2</sup> (n=51) | 3 ± 1 | 3 ± 1 | 2 ± 0.3 | < 0.001 |
| PVR, Wood units (n=50) | 8 ± 5 | 5 ± 2 | 11 ± 6 | < 0.001 |

**Figure 1:**
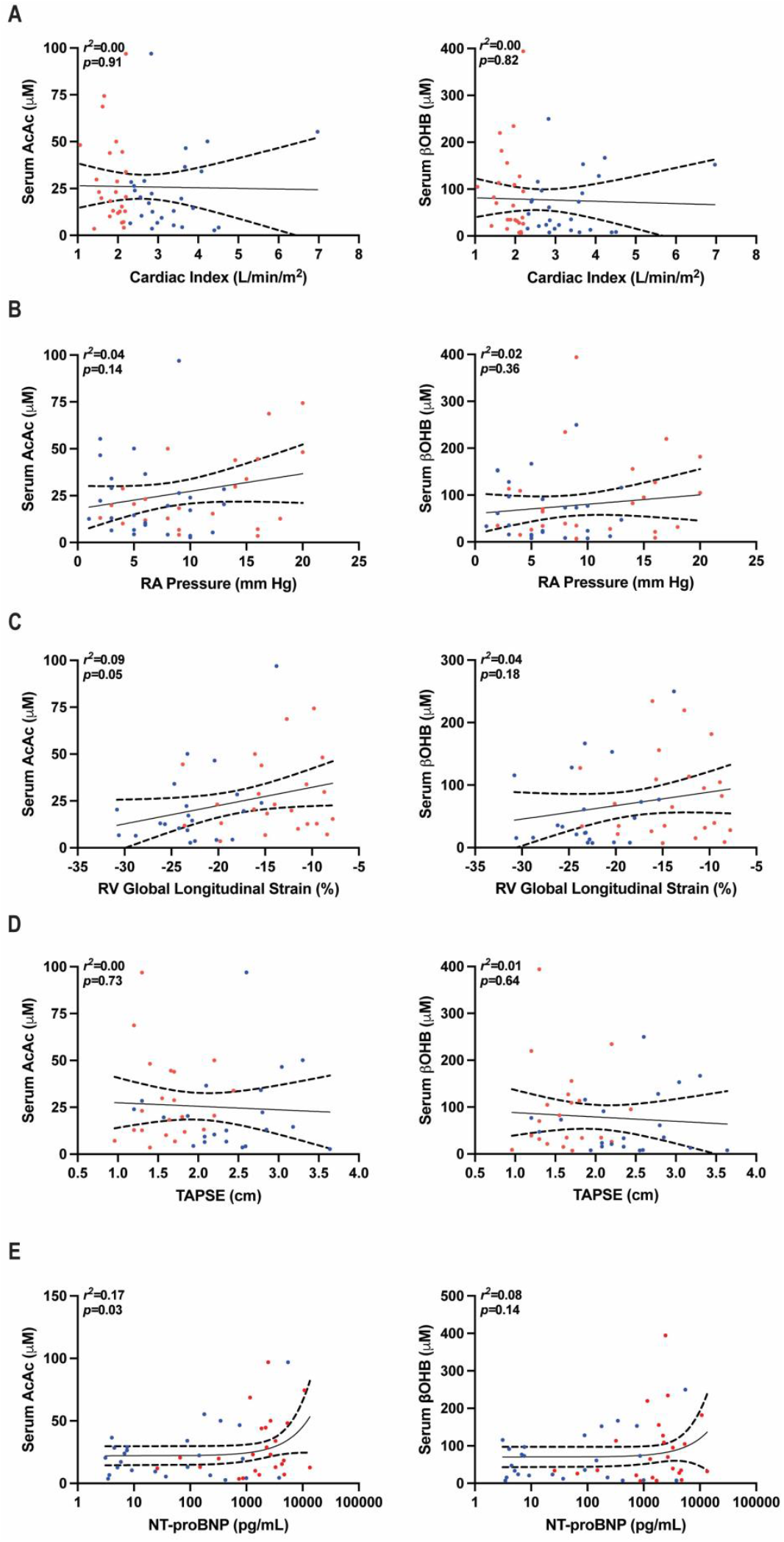
Serum ketones were not associated with echocardiographic, hemodynamic, or biochemical measures of RV function. (A) Cardiac index was not associated with serum ketone body concentrations. (B) Right atrial pressure was not strongly associated with AcAc or βOHB. (C) RV global longitudinal was not associated with serum ketone body concentrations. (D) TAPSE was not significantly associated with AcAc or βOHB. (E) NT-proBNP was not strongly associated with serum ketone body concentrations. Red dots signify decompensated patients (*n*=25) and blue dots signify compensated patients (*n*=26).

Unlike prior observations in patients with LVF (10), circulating concentrations of the two ketone bodies acetoacetate (AcAc) and βOHB were not different when the compensated and decompensated groups were compared (**Supplemental Figure 1**). Surprisingly, AcAc and βOHB levels were not strongly associated with cardiac index, right atrial pressure, RV global longitudinal strain, tricuspid annular plane systolic excursion (TAPSE), or serum NT pro-BNP (**Figure 1**). Additionally, there were no significant relationships between mPAP and PVR and AcAc and βOHB (**Supplemental Figure 2**).

**Figure 2:**
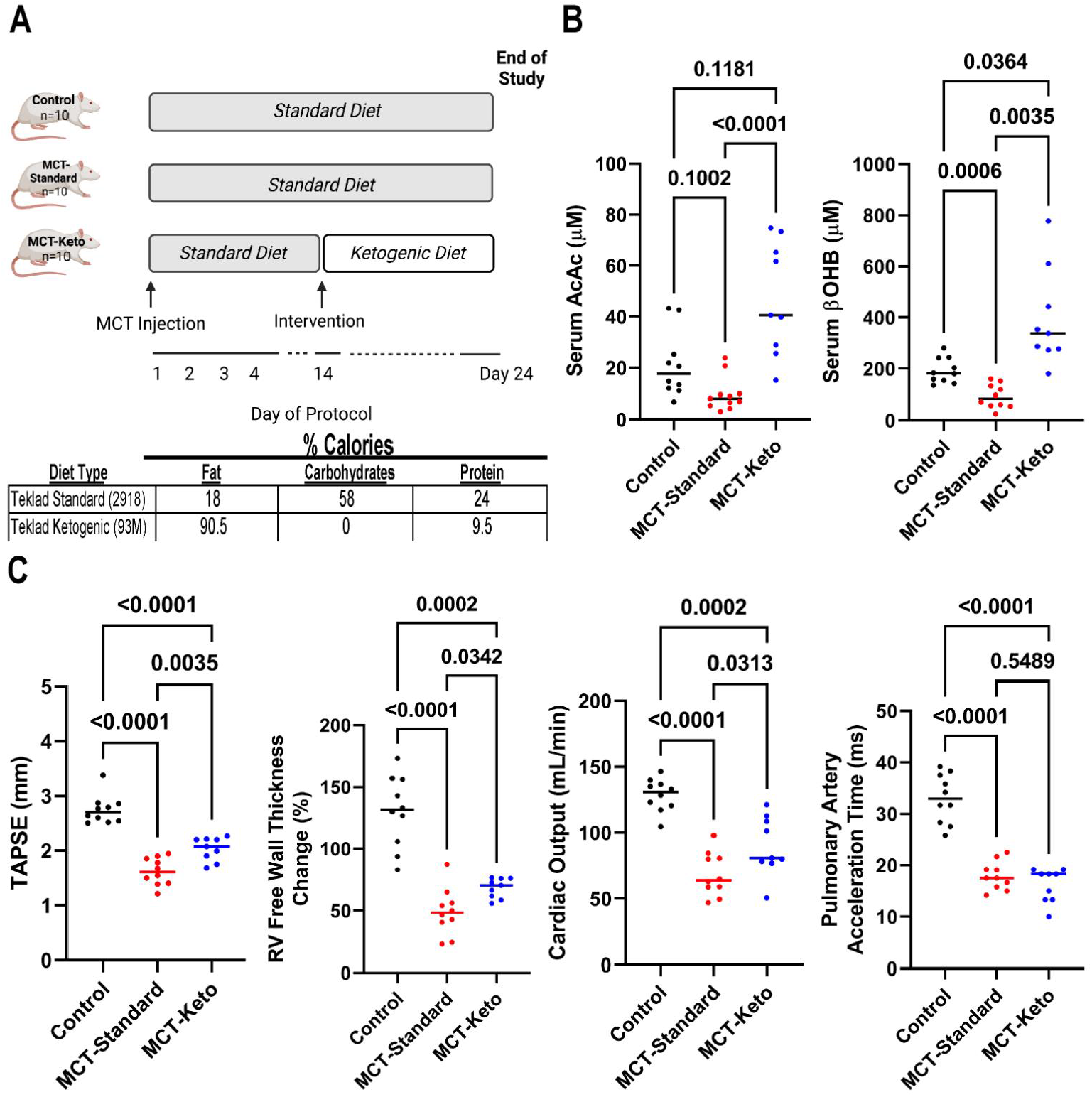
A ketogenic diet increases serum ketone body concentrations and augments RV function in monocrotaline rats. (A) Diagram of experiment approach and caloric composition of diets. (B) A ketogenic diet increased serum concentrations of AcAc and βOHB in MCT-Keto rats. (C) Ketogenic diet significantly increased TAPSE (control: 2.7±0.2, MCT-standard: 1.6±0.2, MCT-keto: 2.0±0.2), RV free wall thickness change (control: 130±29, MCT-standard: 49±19, MCT-keto: 68±8), and cardiac output (control: 129±12, MCT-standard: 65±14, MCT-keto: 90±22) without changing pulmonary artery acceleration time (control: 33±5, MCT-standard: 18±3, MCT-keto: 16±3).

### A Ketogenic Diet Increased Serum Ketone Body Concentrations and Improved RV Function in Rodent RVF

Next, we determined how a dietary-induced ketosis, starting after PAH was allowed to develop, modulated RVF in MCT rats (**Figure 2A**). In agreement with our human data, MCT-Standard rats were not ketotic, and in fact they had lower βOHB levels than controls (**Figure 2B**). Importantly, the ketogenic diet (MCT-Keto) significantly increased serum AcAc and βOHB (**Figure 2B**). Then, we quantified the RV-effects of the ketogenic diet using RV-focused echocardiographic analysis. As compared to MCT-Standard, MCT-Keto rats had higher TAPSE, percent RV free wall thickening, and cardiac output despite no differences in pulmonary artery acceleration time, a well validated echocardiographic marker of pulmonary hypertension severity (**Figure 2C**). Thus, a diet-induced ketosis augmented RV function in preclinical RVF.

### Ketogenic Diet Blunted NLRP3 Inflammasome Activation and Significantly Reduced Macrophage Accumulation in the RV

Finally, we probed how the ketogenic diet intervention modulated RV macrophage infiltration and NLRP3 inflammasome activation. Immunoblots showed MCT-Keto animals had decreased RV levels of NLRP3, pro-caspase-1, pro and mature interleukin-1β, and apoptosis-associated speck-like protein containing a caspase recruitment domain (ASC) relative to the MCT-Standard group, but not all changes were statistically significant (**Figure 3A**). In accordance with our Western blots, confocal microscopy revealed that both total macrophage and ASC+ macrophage abundances in the RV were normalized by the ketogenic diet (**Figure 3B**). Additionally, RV fibrosis was blunted in MCT-Keto animals (**Figure 3C**). In summary, our data showed a ketogenic diet suppressed pathological NLRP3 activation, macrophage infiltration, and RV fibrosis in MCT rats, and these results may explain the RV-enhancing effects the ketogenic diet afforded.

**Figure 3:**
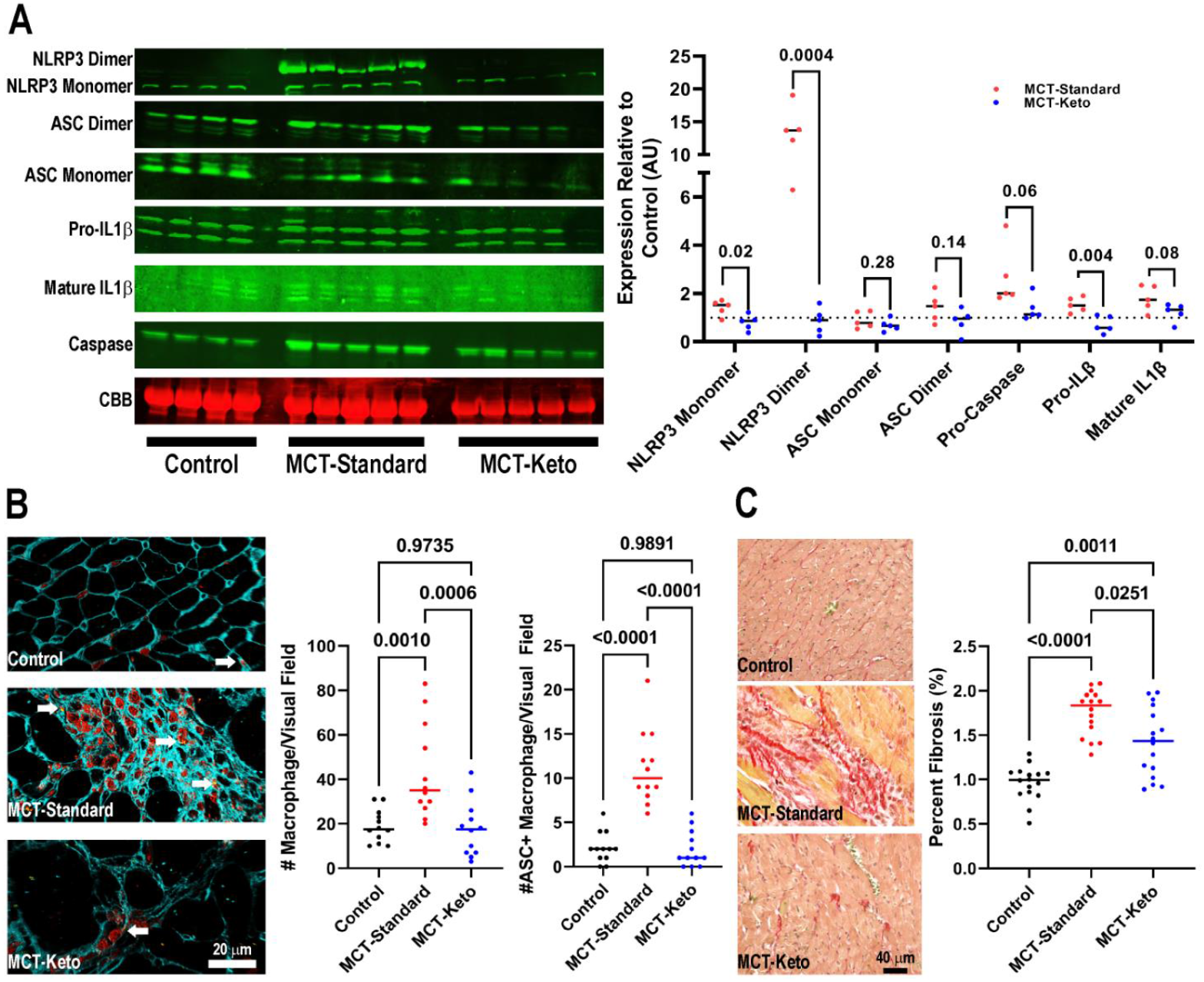
A ketogenic diet suppresses macrophage NLRP3 activation and RV fibrosis in monocrotaline rats. (A) Representative Western blots and subsequent quantification of protein abundance from *n*=4 control, *n*=5 MCT-Standard, and *n*=5 MCT-Keto RV extracts. Signals from the four control animals were averaged to serve as an arbitrary standard of 1. MCT-Standard and MCT-Keto were then compared to each other. (B) Representative confocal micrographs stained with wheat germ agglutinin (Blue), galectin-3 (Red), and ASC (Yellow) to show total macrophage (galectin-3 positive) and ASC+ macrophage in RV sections. Arrows highlight ASC+ macrophage in each section. Quantification of total and ASC+ macrophage in four randomly selected areas per RV section on a 20x objective from three distinct animals per experimental group. (C) Representative Picrosirius Red section with quantification of percent fibrosis on right. MCT-Keto rats had less RV fibrosis than MCT-Standard.

## Discussion

In summary, we show circulating ketone body concentrations are not elevated in human PAH-mediated RVF and they do not change as the severity of RVF, as determined by three distinct but complementary approaches, increases. This is in direct opposition to what is observed in LVF as more severe LV dysfunction results in an even more pronounced ketosis (10). In rodent studies, a ketogenic diet augments RV function and suppresses pathological macrophage NLRP3 inflammasome activation and RV fibrosis. We provide evidence that one method to enhance ketosis improves RV function, potentially through an anti-inflammatory mechanism. Overall, our data suggests ketogenic interventions may have therapeutic relevance for a highly morbid and untreatable form of heart failure.

The absence of ketosis in PAH-induced RVF agrees with other data showing this biochemical response is dysregulated in isolated RVF. In particular, circulating βOHB concentrations in patients with arrhythmogenic right ventricular cardiomyopathy with isolated RV-involvement are lower than those in patients with biventricular involvement (11). Certainly, our data does not dispute the fact that others have documented a ketosis in PAH and chronic thromboembolic pulmonary hypertension (CTEPH), another form of pulmonary hypertension that can compromise RV function. However, the novelty of our data is that the degree of ketosis is not enhanced when RVF is present. Interestingly, Heresi *et al*., also found no significant association between serum βOHB and cardiac index in a cohort of 33 CTEPH patients (12). Finally, Nielsen *et al*., found patients with CTEPH (*n*=10) and PAH (*n*=10) have nearly identical serum βOHB levels, but in this study, the CTEPH patients had significantly worse RV function as determined by strain echocardiography (13). Thus, our data are in line with other groups with smaller sample sizes and the summation of these studies suggests patients with isolated RVF lack a compensatory ketosis.

An additional explanation for the observed benefits afforded by the ketogenic diet in our rodent studies may be the favorable metabolic properties that ketone bodies possess. Ketone bodies are an energetically efficient fuel source, and thus can overcome mitochondrial dysfunction (15). RV failure is characterized by mitochondrial deficits that culminates in the loss of the RV’s ability to utilize its preferred energy source: fatty acids (16-18). It is possible that ketone bodies help the dysfunctional RV overcome the metabolic inflexibility and thereby restore RV bioenergetics. These important metabolic changes could serve as an addition to the anti-inflammatory effects we observed. To gain more insight into this hypothesis, we queried previously published proteomics data, and found 3-hydroxybutyrate dehydrogenase (BDH1), the enzyme that converts βOHB to AcAc and thus serves as the initiating event in ketone oxidation, is actually downregulated in the MCT-RV (**Supplemental Figure 3**). However, 3-oxoacid CoA-transferase 1 (OXCT1), the enzyme that governs ketone catabolism into the tricyclic acid cycle, is unaltered in MCT-RV (**Supplemental Figure 3**) (14). These data suggest that the anti-inflammatory properties of ketones may have been more important than the bioenergetic changes in our preclinical studies. In contrast, proteomics data from human RVF show BDH1 is significantly upregulated while OXCT1 levels are unchanged in the failing RV (**Supplemental Figure 3**) (15). The divergent species response in RV ketone metabolism suggests ketone bodies may have even greater therapeutic utility in human RVF via their synergistic anti-inflammatory and metabolic effects.

At present, approaches that induce ketosis are being investigated in pulmonary hypertension, and these studies are starting to shed light on the utility of ketogenic approaches in RVF. Interestingly, a recent study demonstrated an acute infusion of βOHB exerts beneficial hemodynamic changes in human PAH, and echocardiographic analysis reveals an augmentation of RV function with βOHB. The specific effects of βOHB on the RV versus the pulmonary vasculature are difficult to differentiate as βOHB appears to have pulmonary vasodilatory properties (13), which could underlie the echocardiographic changes in RV function. Nonetheless, this acute study highlights the potential utility of ketosis in PAH, but chronic studies are still needed to evaluate both the long-term therapeutic efficacy and RV-remodeling capabilities. Additionally, a second clinical trial is evaluating how a ketogenic diet impacts symptoms and disease severity in pulmonary hypertension due to heart failure with preserved ejection fraction (NCT04942548). Echocardiography will be used as a secondary endpoint in this study, and hopefully this trial will provide additional insights in the role that ketone bodies play in RV function.

Certainly, future studies are required to gain a deeper understanding of our proposed RV-liver axis, and in particular, a detailed analysis of systemic ketone metabolism in RVF will be crucial. Additionally, it will be important to understand the molecular consequences of RVF on hepatic metabolic function. Certainly, there are clear clinical observations that chronic RVF can manifest as cirrhosis, but the mechanisms underlying this phenomenon are unknown. Defining the molecular mediators of RVF-mediated hepatic dysfunction are sorely needed as these studies may illuminate therapeutic targets for this highly morbid complication of RVF (4).

Finally, our study has important limitations that we must acknowledge. First, while the ketogenic diet increases AcAc and βOHB in preclinical PAH, we cannot be rule out that the beneficial effects may be due to low carbohydrate or high fat content. Second, our rodent intervention only involved male rats because they develop more severe RV failure than females. Potential sex-specific differences in response to increased ketone levels were not the aim of this study, but they should be evaluated in the future.

## Methods

### Human Patients

We examined the relationship between RV function and serum ketone bodies in 51 PAH-patients from the University of Minnesota Pulmonary Hypertension Program. PAH patients were classified into two groups: a compensated (cardiac index>2.2 L/min/m^2^ as defined by Thermodilution) or a decompensated (cardiac index ≤2.2 L/min/m^2^) RV phenotype (**Table 1**). Patients with known hepatic conditions were excluded. RV function was characterized using hemodynamic and comprehensive echocardiographic analysis (RV strain, RV free wall arterial compliance (RVFAC), RV end diastolic area (RVEDA), RV end systolic area (RVESA), tricuspid annular plane systolic excursion (TAPSE), S’, tricuspid regurgitation severity) (**Table 1**). Echocardiography images were analyzed offline and blindly by FK. Human studies were approved by the University of Minnesota Institutional Review Board.

### Serum Ketone Measurements

Fasting blood samples were drawn from each patient prior to a planned hemodynamic assessment. Circulating serum ketone bodies acetoacetate (AcAc) and beta-hydroxybutyrate (βOHB) were quantified using ultra performance liquid chromatography and mass spectrometry (UPLC-MS/MS) (16).

### Rodent Ketogenic Intervention

Male Sprague Dawley rats (Charles River Laboratories) were randomized into three experimental groups: phosphate buffered saline injected (control), monocrotaline (MCT, 60 mg/kg subcutaneous injection) rats fed standard (MCT-Standard) chow (Teklad:2918), and monocrotaline rats fed ketogenic (MCT-Keto) chow (Teklad:93M). To ensure a translational approach, dietary intervention began two weeks after MCT injection. End point analyses were performed 24 days post-MCT exposure. Rodent RV function was examined with echocardiography using a Vevo2100 ultrasound system as previously described (17). The University of Minnesota Institutional Animal Care and Use Committee approved rodent studies.

### Serum Ketone Measurements

Animals were fasted overnight and then blood was drawn for analysis. Blood was allowed to coagulate for 30 minutes and then was spun at 1000*g* for 20 minutes; the serum fraction was aspirated, and immediately snap frozen in liquid nitrogen. Ketone bodies were measured using UPLC-MS/MS as described above.

### RV Fibrosis Assessment

RV free wall sections were collected and fixed in 10% formalin before being embedded in paraffin, sectioned at 10-μm, and stained with Picrosirius Red by the University of Minnesota Histology and Research Laboratory in the Clinical and Translational Science Institute.

### Confocal Microscopy

RV free wall sections were deparaffinized using Xylene and subsequent incubations with 100% ethanol, 95% ethanol, and 70% ethanol. Slides were then placed in a water bath with 10% decloaking solution, before being permeabilized with 1% Triton X-100 in PBS. Slides were blocked with 5% goat serum before incubation with primary antibodies galectin-3 and ASC overnight at 4°C. Following primary antibody incubation, sections were blocked with 5% goat serum before incubation with VectaFluor Amplifier Antibody. Secondary antibody incubation was performed for 30 minutes at 37°C. Sections were washed with PBS and incubated with 0.1% Hoechst stain before treatment with a fluorescence quenching kit and mounting in anti-fade reagent. Confocal micrographs were collected on a Zeiss LSM 900 Airyscan 2.0 microscope. RV fibrosis and macrophage analyses were blindly performed by MB.

### Immunoblots

RV free wall extracts were prepared and subjected to Western blotting protocol to quantify protein expression as previously described (18). Antibodies to NLRP3, Caspase-1, interleukin-1β, and apoptosis-associated speck-like protein containing a CARD (ASC) were used to biochemically characterize inflammasome activation. A complete list of antibodies used for immunoblots is included in **Supplemental Table 1**.

### Statistical Analysis

All data were evaluated for normality using Sharpio-Wilk test. When analyzing differences between two groups, normally distributed variables were compared using unpaired *t*-test and Mann-Whitney U-test was used for non-normally distributed variables. When comparing three groups with normal distributions, one-way ANOVA with Tukey post hoc analysis was performed if variance was equal as determined by Brown-Forsythe test. If variance was unequal, Brown-Forsythe and Welch ANOVA with Dunn post hoc analysis was completed. If the data were not normally distributed, the Kruskal-Wallis and Dunn’s multiple-comparisons tests were employed. Linear regression analysis was used to determine the relationships among serum ketone concentrations, RV function, and PAH severity. All statistical analyses were performed on GraphPad Prism version 9.0. Statistical significance was defined by *p*<0.05.

---

**Supplemental Figure 1:**
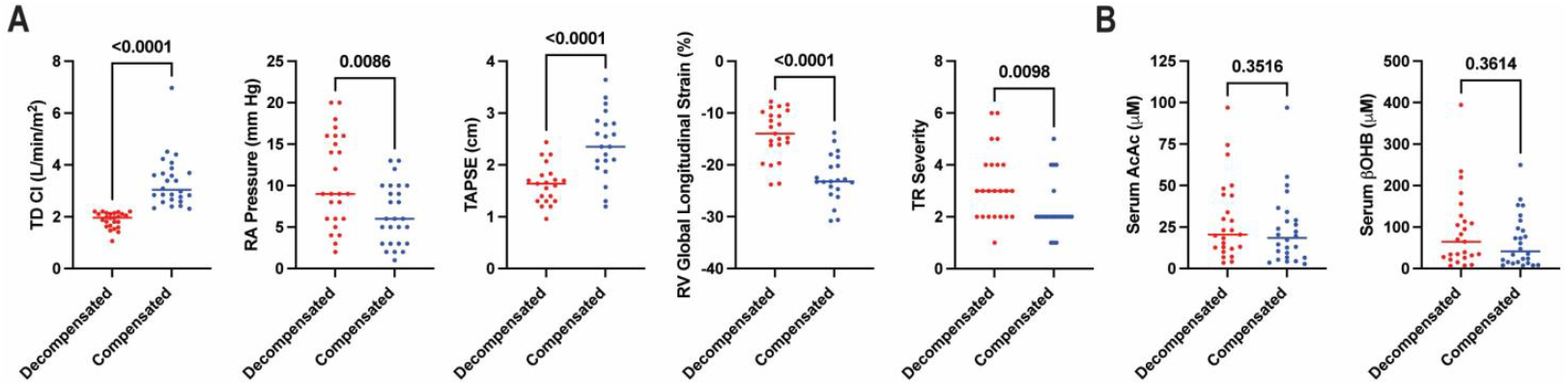
Serum ketones were not elevated in PAH patients with RV dysfunction. (A) Both hemodynamics and echocardiography revealed a greater degree of RV dysfunction and tricuspid regurgitation severity in the decompensated group. (B) Serum AcAc and βOHB concentrations were not elevated in the decompensated group.

**Supplemental Figure 2:**
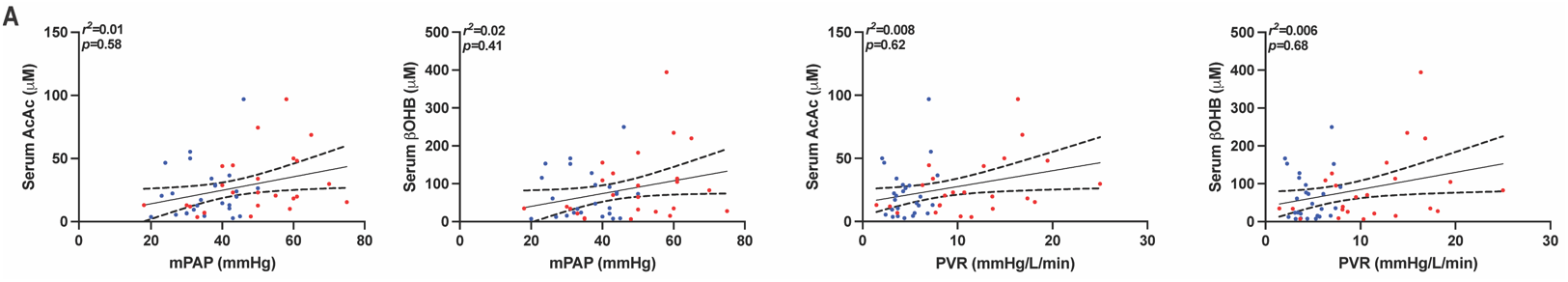
Serum ketones were not associated with pulmonary vascular disease severity. (A) Hemodynamic measurements of PAH severity were not associated with serum ketone body concentrations.

**Supplemental Figure 3:**
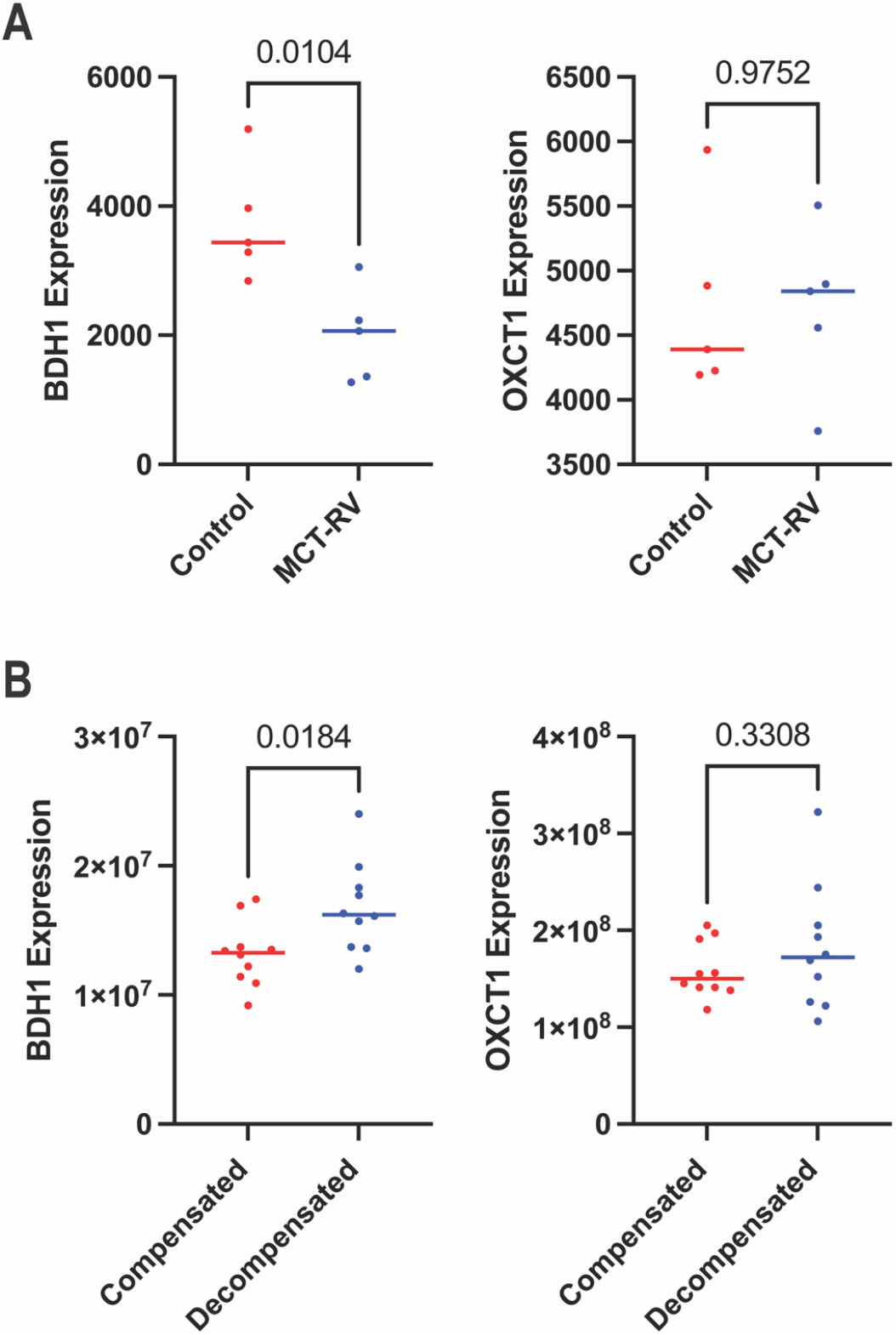
The ketolytic enzymes BDH1 and OXCT1 exhibit divergent responses in rodents and humans with RVF. (A) BDH1 is significantly downregulated, but OXCT1 levels are unchanged in the MCT RV. (B) BDH1 levels are significantly higher and OXCT1 abundance is unchanged in human RVF.

**Supplemental Table 1:** Antibodies/Fluorophores Used for Immunoblots and Confocal Microscopy.

| Antigen | Company | Species | Dilution |
| --- | --- | --- | --- |
| **Western Blots** |  |  |  |
| Caspase-1 | Abcam | Rabbit | 1:250 |
| NLRP3 | Novus | Rabbit | 1:250 |
| IL-1B | Abcam | Rabbit | 1:250 |
| ASC | Abcam | Rabbit | 1:250 |
| Anti-mouse 680 | Li-COR | Goat | 1:5000 |
| Anti-rabbit 800 | Li-COR | Goat | 1:5000 |
| **Immunofluorescence** |  |  |  |
| Galectin-3 | Abcam | Rabbit | 1:50 |
| ASC | Abcam | Goat | 1:50 |
| Anti-rabbit Alexa Fluor 568 | Invitrogen | Goat | 1:500 |
| Anti-goat Alexa Fluor 647 | Thermo Scientific | Donkey | 1:500 |
| Wheat Germ Agglutinin Alexa Fluor 633 | Invitrogen |  | 1:50 |

